## Supplemental files for "Multiple polarity kinases inhibit phase separation of F-BAR protein Cdc15 and antagonize cytokinetic ring assembly in fission yeast"

**Supplemental Figures.**

**Figure 1- figure supplement 1. In vitro kinase assays to show Cdc15C is directly phosphorylated on its C-terminus by Kin1, Shk1 or Pck1.** (A) Kin1, Shk1 and Pck1 directly phosphorylate MBP-tagged Cdc15C whereas the N-terminus fragment of Cdc15 (MBP-Cdc15(N)) remains unphosphorylated. (B) Mutation of Pom1 phosphorylation sites on GST-tagged Cdc15C to alanine (Ala) or aspartic acid (Asp) does not prevent its phosphorylation by Shk1, Kin1, or Pck1. Kinase reactions were analyzed by SDS-PAGE, stained with Coomassie Blue (CB) and $^{32}P$ incorporation was detected by autoradiography.

**Figure 3- figure supplement 1,2 and 3. Mapping of phosphorylation sites of Kin1, Shk1 and Pck1 on Cdc15.** (A) Representative mass spectra of phosphopeptides from Cdc15 *in vitro* phosphorylated by Kin1 (figure supplement 1), Shk1 (figure supplement 2), or Pck1 (figure supplement 3) are shown. Images were extracted from Scaffold PTM; matched b and y ions are highlighted in red and blue, respectively. Ions resulting from neutral losses are highlighted in green. When available, both MS2 and MS3 (spectrum resulting from further fragmentation of the neutral loss (NL) of the phosphate from the parent ion during MS2 scan) spectra of the same site are shown. Sp/Tp: phosphorylated serine/threonine; Sd/Td: dehydrated serine/threonine (due to neutral loss of phosphate). (B) GST tagged Cdc15C was phosphorylated by MBP-Kin1, GST-Shk1 and MBP-Pck1 in presence of $\gamma$-[$^{32}P$]-ATP for 30 minutes. Proteins were digested with trypsin and peptides were separated by thin-layer electrophoresis and chromatography. Phosphopeptides were detected by autoradiography.

**Figure 5- figure supplement 1. mNG-Cdc15 localization after treatment with DMSO or inhibitor(s) and in Cdc15 phosphomutants.** (A) Representative images of the indicated strains (also shown in Fig. 5A) to demonstrate mNG-Cdc15 localization after treatment with inhibitor(s). All representative images were deconvolved and max projected using Image J. (B) Representative images of Figure 5B showing membrane-cytosol localization of the indicated Cdc15 phosphomutants.

**Figure 5- figure supplement 2. Quantification of tip-septa of the indicated strains** (A) Quantification of the tip septa phenotype of the indicated strains after treatment with DMSO or the inhibitor(s) 3-MB-PP1 or 3-BrB-PP1 for 2 hr at 32°C from 3 biological replicates. N≥ 100 cells.

**Figure 6- figure supplement 1. Comparison of HPS and KH D models for Cdc15 dimers and convergence properties of simulations.** (A, B) Snapshots of initial configurations for both the HPS and KH D models. Ten independent simulations of the Cdc15 dimer in each phosphorylation state were performed of at least 1.9 μs, with half initialized with the IDRs apart and half with the IDRs interacting. A higher temperature of 350K was used for KH D model. (C) Representative simulation snapshots at the indicated COM distance between each IDR. (D) Frequency distribution of distance between the COM of each IDR in the KH D model, averaged over all simulations (as in Figure 6F for HPS model). Grey line indicates 100 Å. A tendency for increased IDR separation with increased phosphorylation similar to the HPS model can be observed. However, as shown in panels E and F, the IDRs mostly remain trapped into fused or segregated states according to their initial conditions of panel A so the bimodal distribution reflects the initial conditions rather than equilibrium. (E).

Frequency distributions of distance between the COM of each IDR in the HPS models, according to the initial condition of panel A. Distributions approach a common curve independent of the initial condition, indicating that simulations approach equilibrium, though less so for the Full and Shk1 cases. (F) Same as panel E, for the KH D model. The COM remains close to the respective initial condition, indicating lack of KH D model equilibration. A fraction of the Pom1 and Full cases however still separate to COM above 100 Å, even if started together.

**Figure 6- figure supplement 2. Cdc15 dimer simulation contact maps in HPS model.** (A, B) Residue to residue contact frequency for the dephosphorylated (Deph) and fully phosphorylated (Full) protein models, calculated with a 10 Å cutoff, over all ten simulations of Figure 6 and Figure 6-figure supplement 1E. (C) Difference of the plots in panels A,B such that blue indicates an increase in contacts in the fully phosphorylated (Full) state. Blue and red striped patterning occurs for the contacts between F-BAR and IDR, and between F-BAR and SH3 domains. The predominantly blue regions, indicating increased contacts, occur at the poles of the F-BAR, while the predominantly red regions occur towards the middle of the F-BAR (see Figure 5-figure supplement 3). The region of inter-chain IDR contacts is red overall, indicating decreased interaction between the IDRs in the fully phosphorylated case.

**Figure 6- figure supplement 3. Cdc15 dimer simulation F-BAR contacts in HPS model.** (A) Spatial depiction the contact frequency of the residues on either side of the F-BAR with any residues in the IDR (sum of contact frequency with residues 301-869 in chain A, or 299-869 in chain B), calculated with a 10 Å cutoff. (B) Same as panel A but for contacts with any residues in the SH3 (sum of contact frequency with residues 870-927 in either chain). The value is averaged over all ten simulations of Figure 6.

**Figure 6- figure supplement 4. Cdc15 IDR size in single chain simulations.** (A, C) $R_G$ vs. temperature in the indicated model. (B, D) Increase in $R_G$ calculated relative to the dephosphorylated (Deph) case in the indicated model. For each model, independent simulations were run at 16 different temperatures chosen to span the coil-to-globule transition. For HPS, trajectory durations for each temperature range from 3.8 to 21.4 μs depending on convergence efficiency, with the most collapsed simulations running for the shortest time. KH D trajectories range from 5.8 to 30.7 μs.

**Figure 6- figure supplement 5. Diagram of States for Cdc15 IDR.** Diagram of states for the Cdc15 IDR (residues 327-854) treating phosphorylation charge as -2e as in the HPS model of Perdikari et al. and in our implementation of KH D model.

**Figure 7- figure supplement 1. Cdc15-IDR-SH3 purification.** (A) Purified Cdc15-IDR-SH3 construct used for in vitro phase separation assays was by resolved by SDS-PAGE and then stained with CB. (B) Before the in vitro droplet assays, purified Cdc15-IDR-SH3 with or without treatment of λ phosphatase (λ) were separated by SDS-PAGE and immunoblotted for Cdc15.

**Figure 8- figure supplement 1. Purification of full length Flag-Cdc15 (E30K, E152K) and His-Fic1.** Purified recombinant proteins used for in vitro droplet assay in Figure 7, (A) Flag-Cdc15 (E30K, E152K) and (B) His-Fic1.

**Figure 8- figure supplement 2. Control images from in-vitro droplet experiments with the purified recombinant proteins.** (A-D) Indicated recombinant proteins displaying no visible droplets in 50 mM Tris pH 7.4, 150 mM NaCl (A,B) or 250 mM NaCl (C,D) with 5% PEG in presence (A) or absence of λ (B-D).

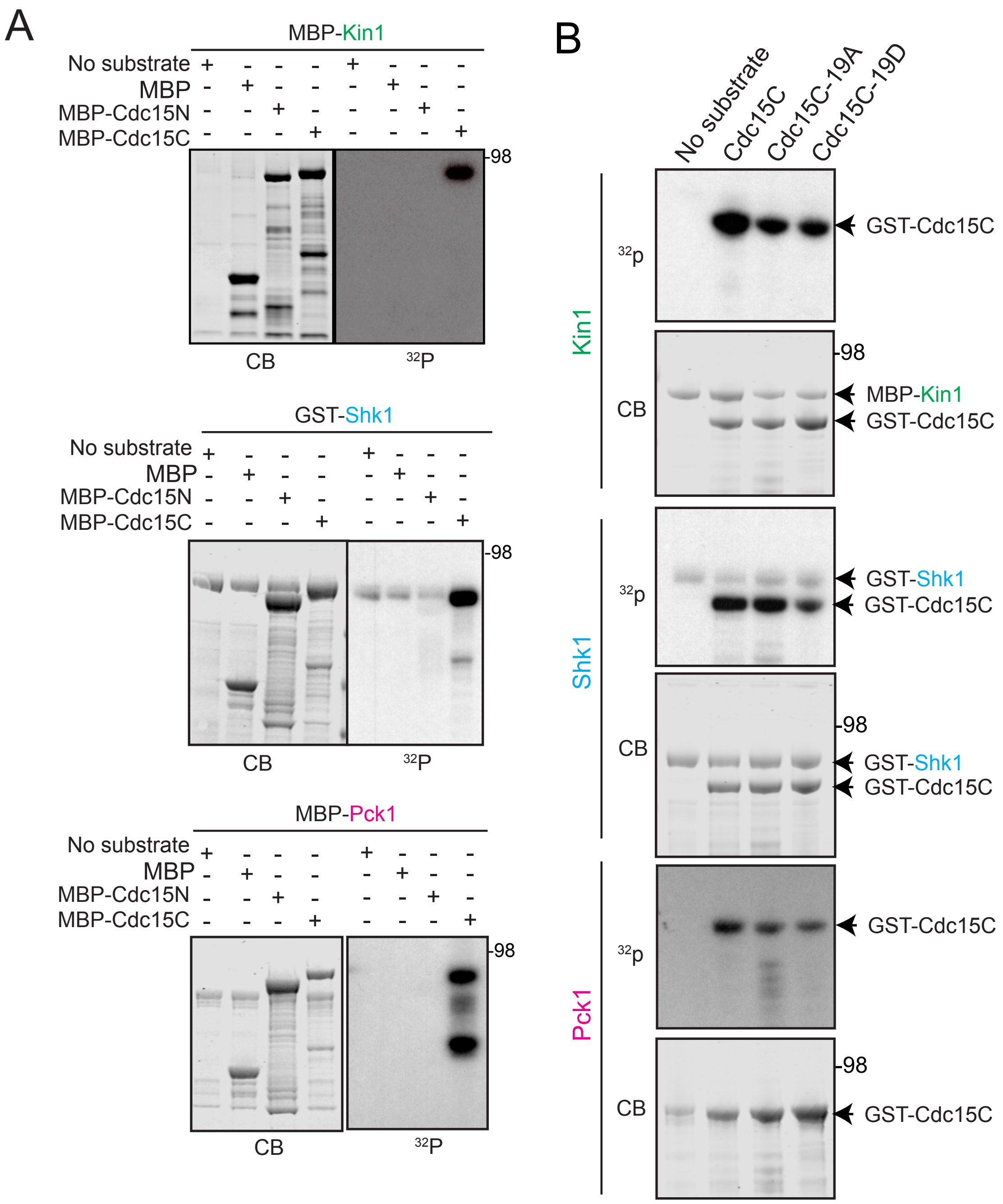


Figure1-figure supplement 1

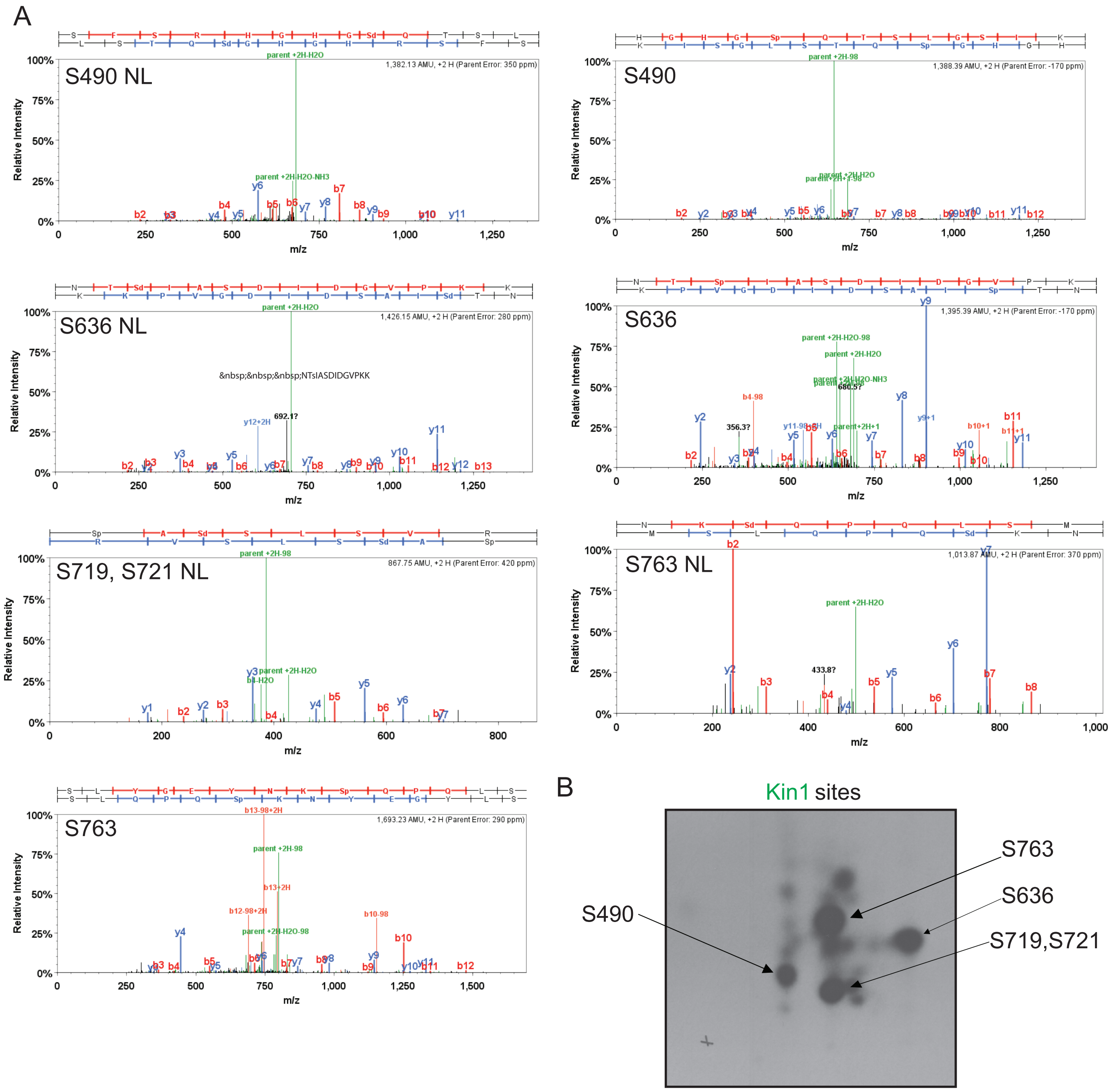


Figure 3- figure supplemental 1

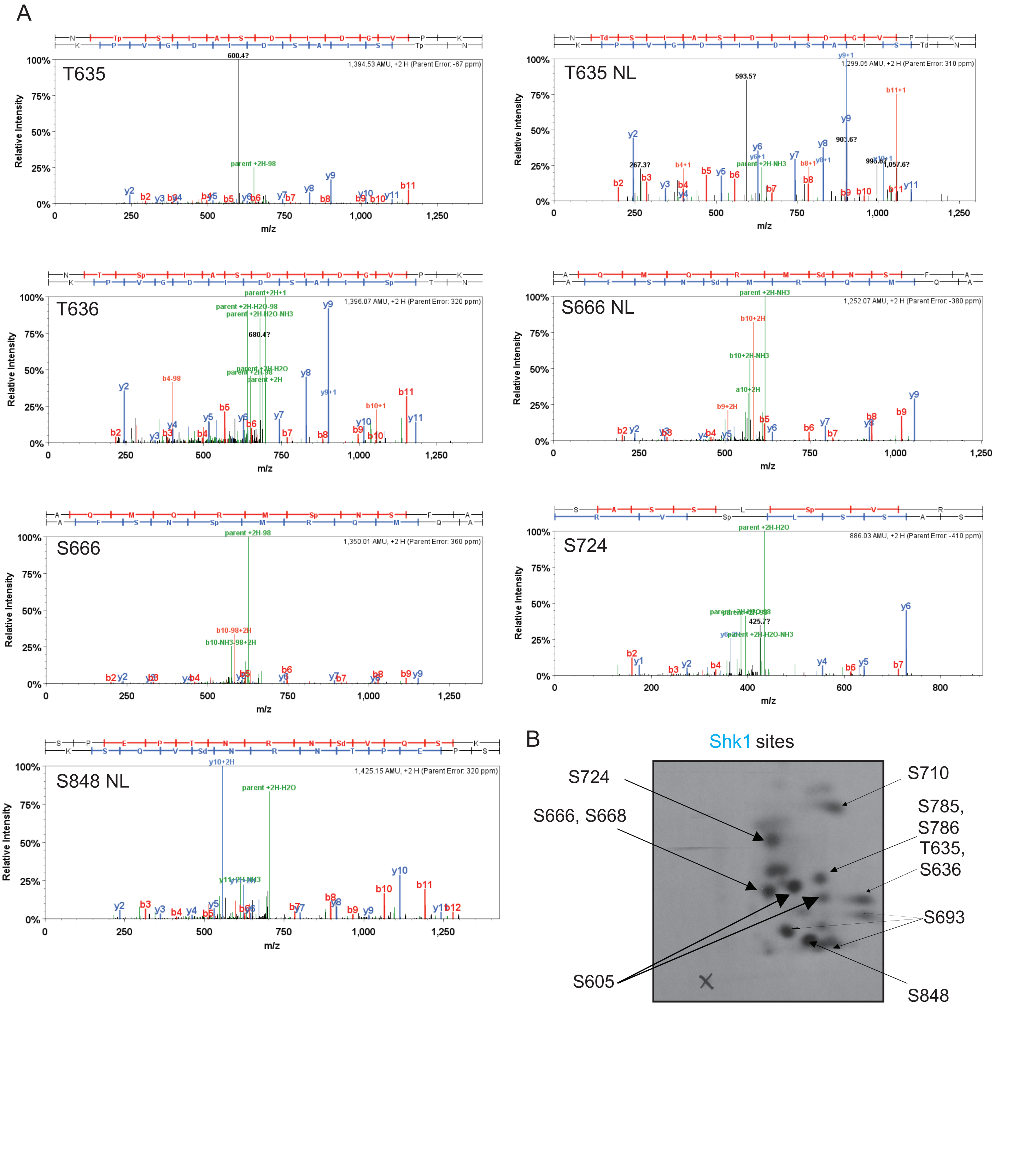


Figure 3- figure supplemental 2

A

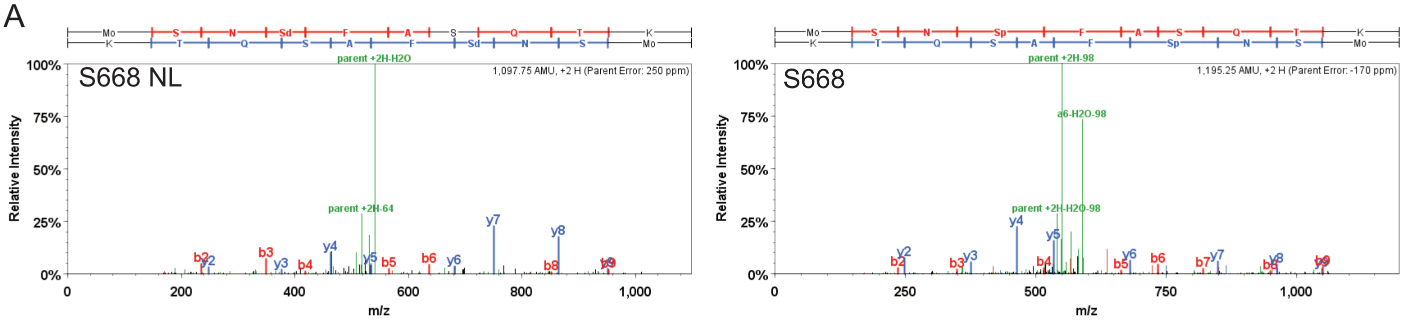


B

Pck1 sites

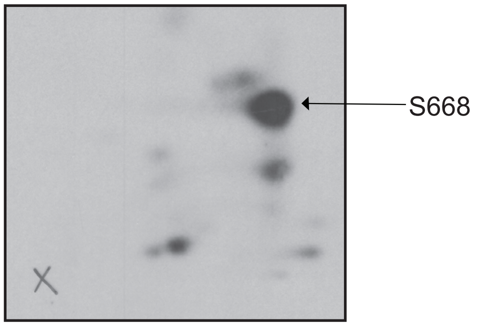


Figure 3- figure supplemental 3

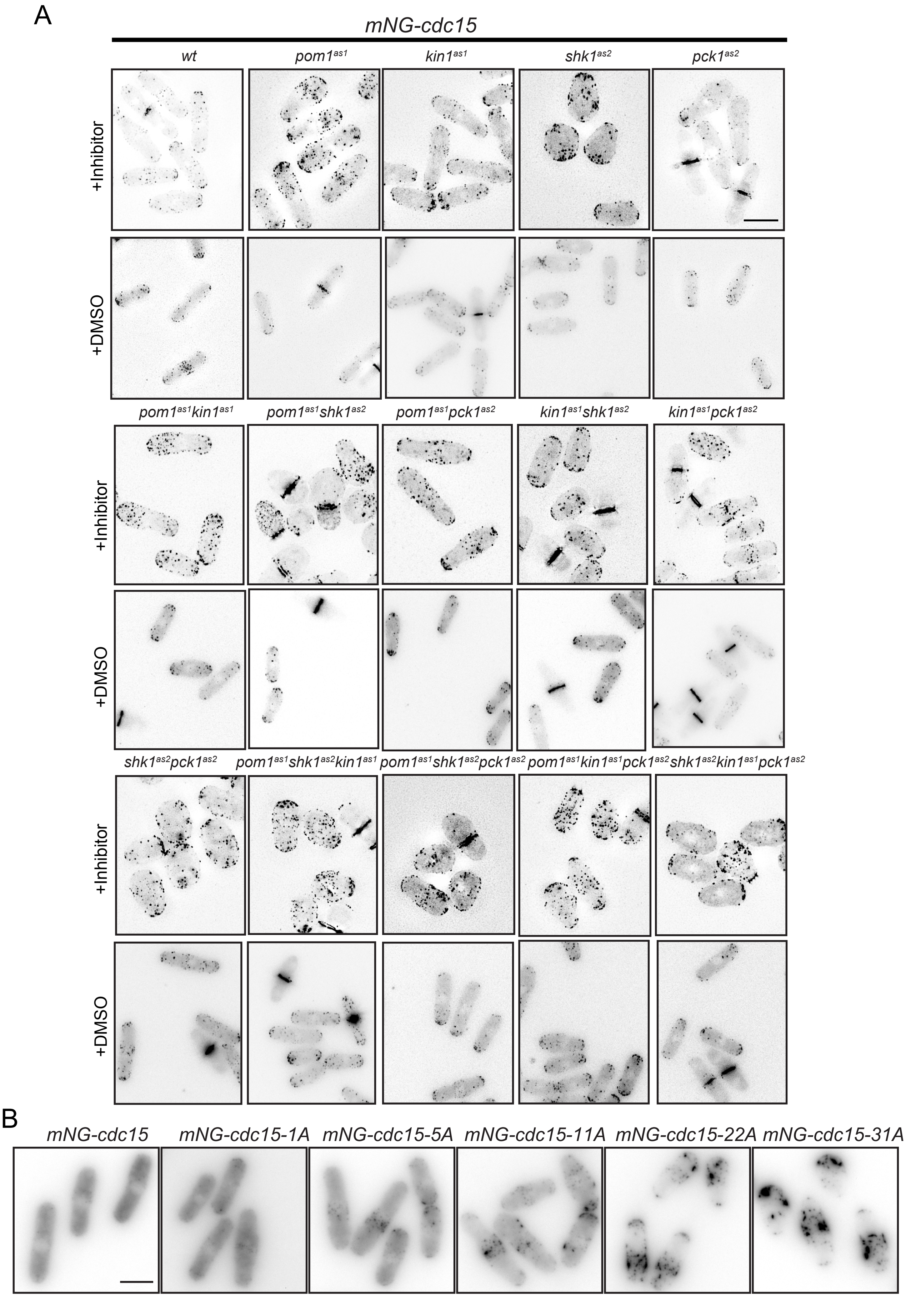


Figure 5-figure supplement 1

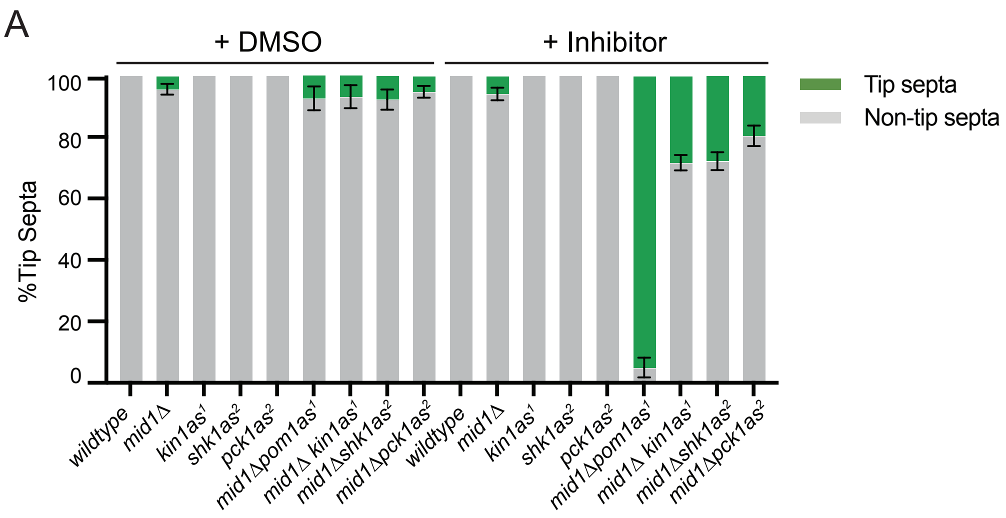


Figure 5-figure supplement 2

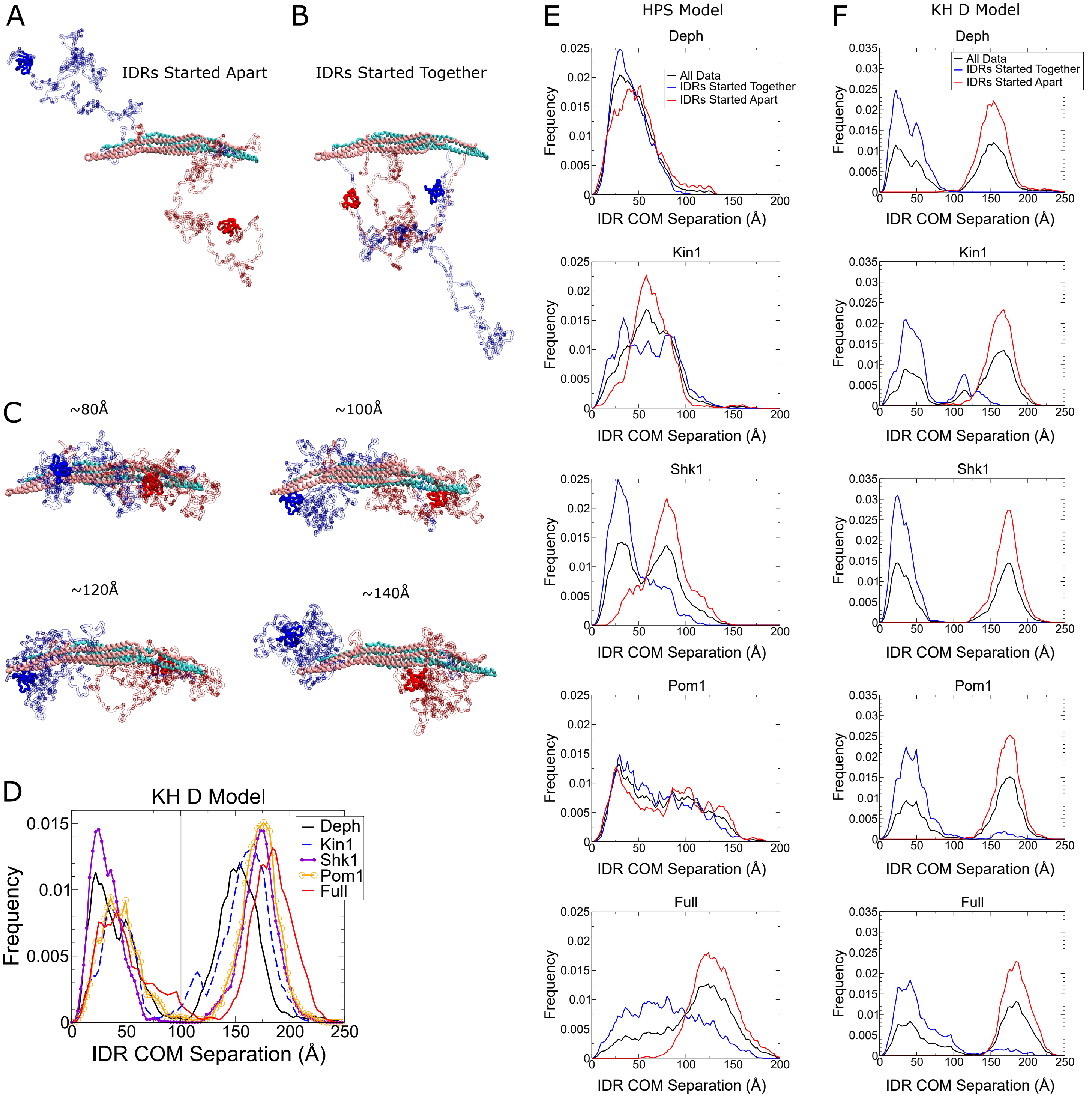


Figure 6-figure supplement 1

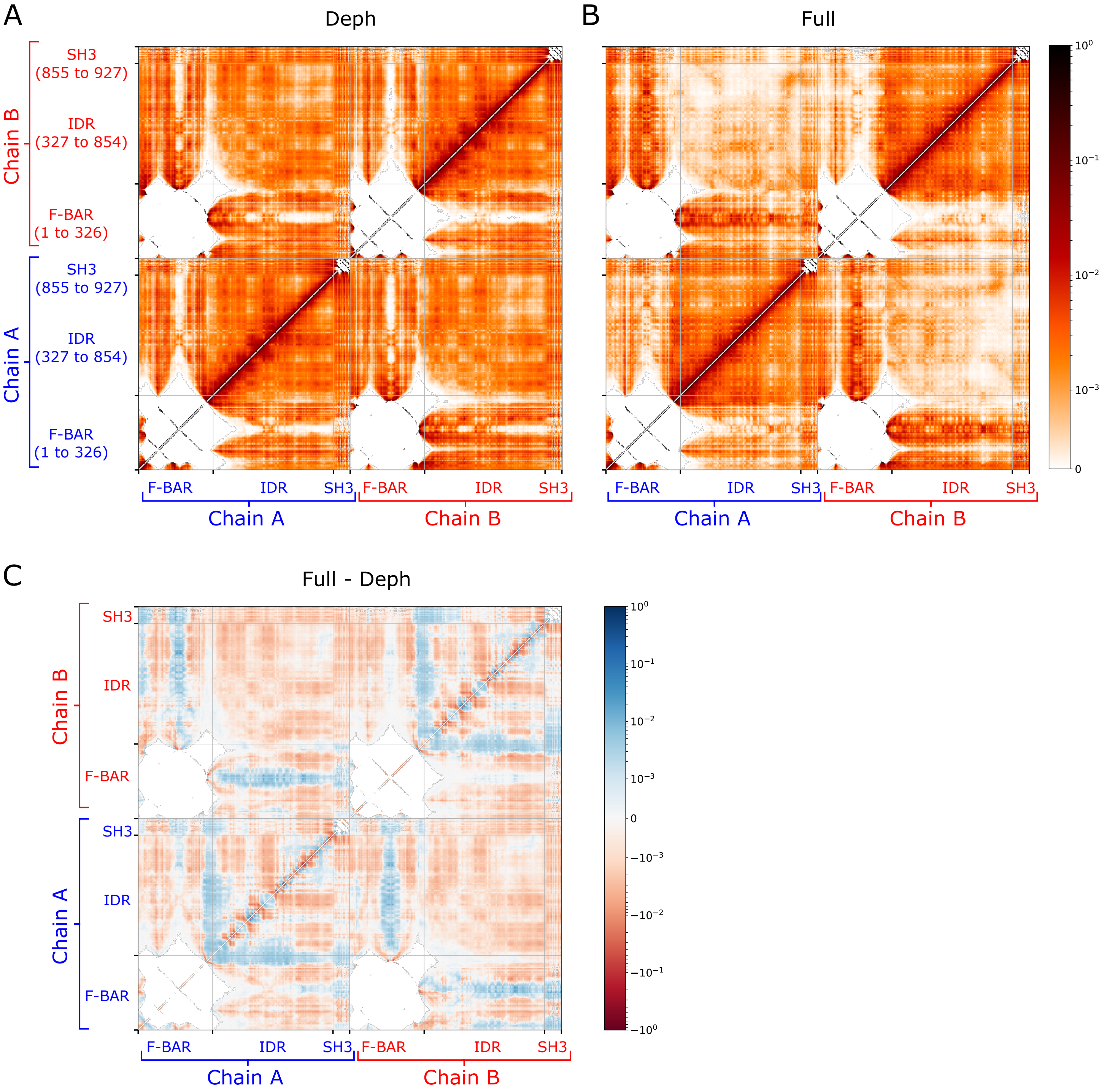


Figure 6-figure supplement 2

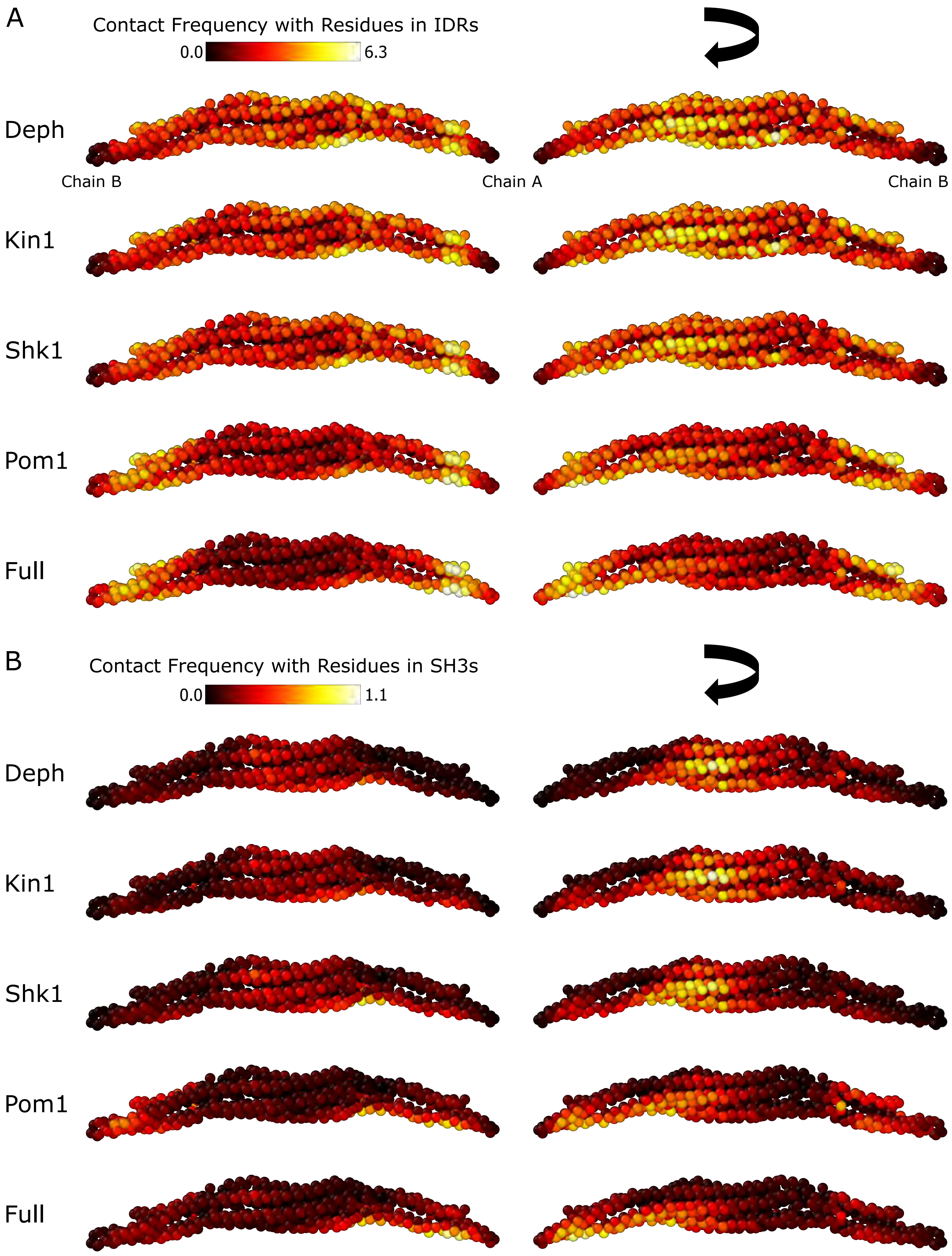


Figure 6-figure supplement 3

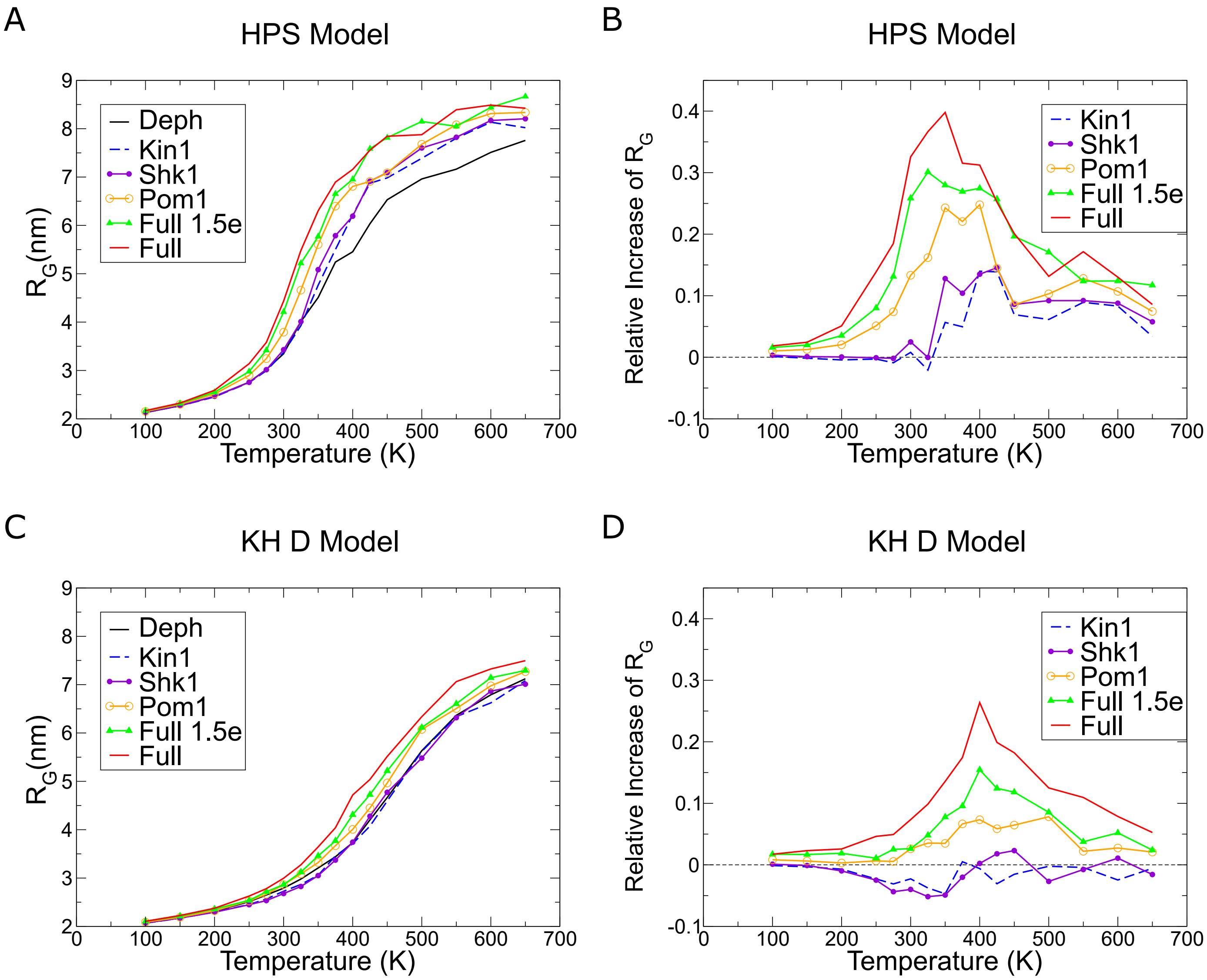


Figure 6-figure supplement 4

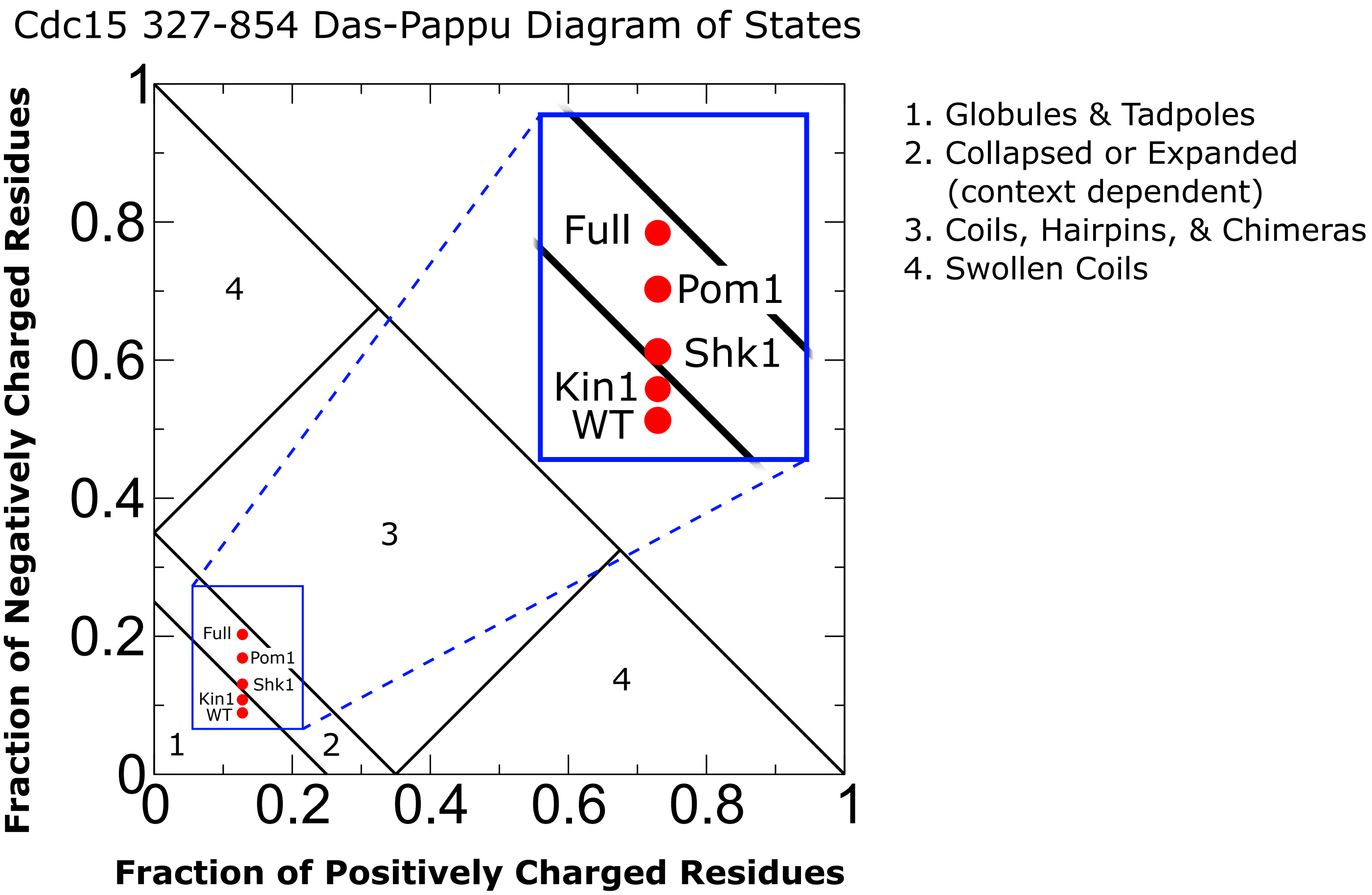


Figure 6-figure supplement 5

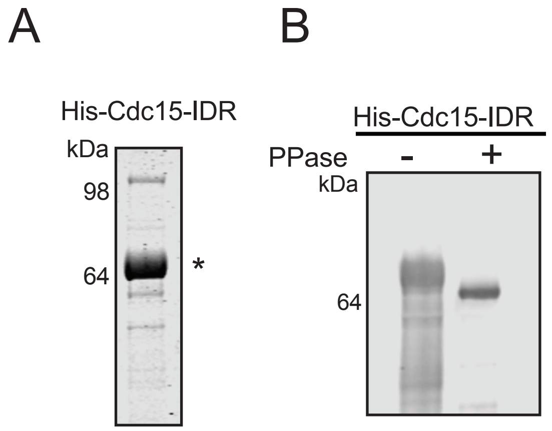


Figure 7- figure supplement 1

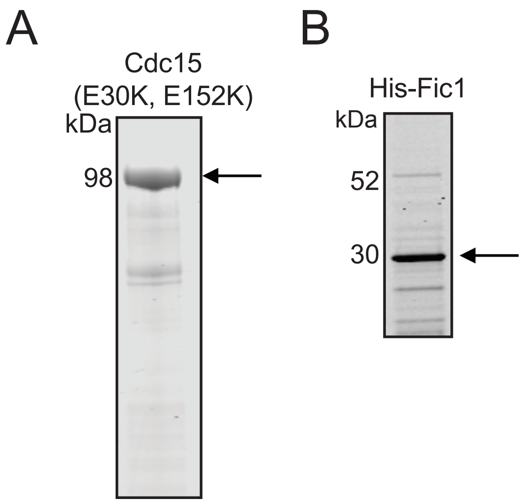


Figure 8- figure supplement 1

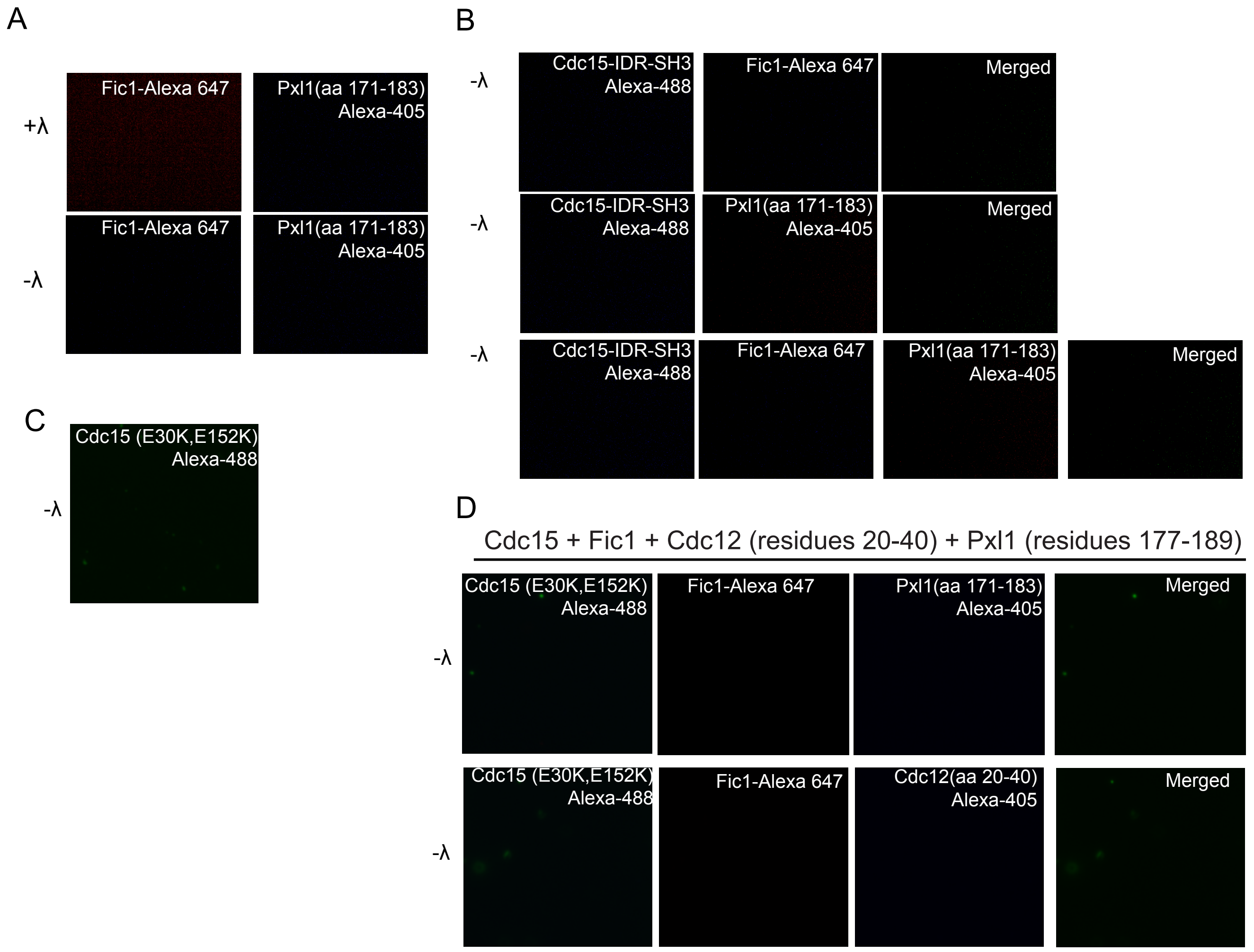


Figure 8-figure supplement 2

### Supplementary Table 1. *S. pombe* strains used in this study

| Strain number | Genotype | Source |
|---|---|---|
| **Figure 1** | | |
| KGY246 | *ade6-M210 ura4-D18 leu1-32 h⁻* | Lab stock |
| KGY11562 | *cdc15-22A ade6-M210 ura4-D18 leu1-32 h⁻* | Lab stock |
| KGY2001-2 | *pom1$^{as1}$-tdTomato:nat$^{R}$ ade6-M210 ura4-D18 leu1-32 h⁻* | (Martin and Berthelot-Grosjean, 2009) |
| KGY1516-2 | *kin1$^{as1}$-FLAG:kan$^{R}$ ura4-294 leu1-32 ade6-704 h$^{+}$* | This study |
| KGY2002-2 | *kin1$^{as1}$-FLAG:kan$^{R}$ pom1$^{as1}$-tdTom:natR leu1-32 ade6-X ura4-X h$^{+}$* | This study |
| KGY15733-2 | *kin1::ura4$^{+}$ ade6-M210 ura4-D18 leu1-32 h$^{+}$* | (Bimbo et al., 2005) |
| KGY11570-2 | *cdc15-22A kin1::ura4$^{+}$ ade6-M21X ura4-D18 leu1-32h-* | This study |
| KGY19282 | *shk1::ClonNat$^{R}$ shk1$^{as2}$(M460A)-Hph$^{R}$ leu1-32 ura4-D18 ade6-M210 h⁻* | (Cipak et al., 2011) |
| KGY1954-2 | *cdc15-22A shk1::ClonNat$^{R}$ shk1-as$^{2}$(M460A)-Hph$^{R}$ ade6-M21X leu1-32 ura4-D18 h⁻* | This study |
| KGY6404 | *pck1::ura4$^{+}$ ade6-M21X ura4-D18 leu1-32h⁻* | This study |
| KGY19615 | *cdc15-22A orb2-34 ade6-M21X leu1-32 ura4-D18 h⁻* | Lab stock |
| KGY1956-3 | *kin1$^{as1}$:kan$^{R}$ shk1::ClonNat$^{R}$ shk1$^{as2}$(M460A)-Hph$^{R}$ pom1$^{as1}$:nat$^{R}$ ade6-M21X leu1-32 ura4-D18 h⁻* | This study |
| KGY1953-2 | *kin1$^{as1}$:kan$^{R}$ shk1::ClonNat$^{R}$ shk1$^{as2}$(M460A)-Hph$^{R}$ ade6-M21X leu1-32 ura4-D18 h⁻* | This study |
| KGY1912-2 | *shk1::ClonNat$^{R}$ shk1$^{as2}$(M460A)-Hph$^{R}$ pom1$^{as1}$:nat$^{R}$ ade6-M21X leu1-32 ura4-D18 h⁻* | This study |
| KGY7477 | *orb2-34 ade6-M21X leu1-32 ura4-D18 h⁻* | (Verde et al., 1995) |
| KGY1962-2 | *kin1$^{as1}$:kan$^{R}$ shk1::ClonNat$^{R}$ shk1$^{as2}$(M460A)-Hph$^{R}$ pom1$^{as1}$:nat$^{R}$ pck1$^{as2}$-HAx3:hyg$^{R}$ ade6-M21X leu1-32 ura4-D18 h⁻* | This study |
| KGY526-2 | *pck1$^{as2}$-FLAG:kan$^{R}$ ade6-M21X leu1-32 ura4-D18 h$^{+}$* | Lab stock |
| KGY19598 | *cdc15-22A pck1$^{as2}$-HAx3:hyg$^{R}$ ade6-M21X leu1-32 ura4-D18 h⁻* | This study |
| **Figure 3** | | |
| KGY246 | *ade6-M210 ura4-D18 leu1-32 h⁻* | Lab stock |
| KGY6012-2 | *cdc15-31A ade6-M210 ura4-D18 leu1-32 h⁻* | This study |
| KGY56 | *nda3-km311leu1-32h$^{+}$* | Lab stock |
| KGY5099 | *cps1-191 leu1-32lys1-151ura4-D18h$^{+}$* | Lab stock |
| **Figure 4** | | |
| KGY5365-2 | *mNG-cdc15 rlc1-mCherry:nat$^{R}$ sid4-mCherry:hyg$^{R}$ leu1-32 ura4-D18 ade6-M210 h⁻* | Lab stock |
| KGY5366-2 | *mNG-cdc15-31A rlc1-mCherry:nat$^{R}$ sid4-mCherry:hyg$^{R}$ leu1-32 ura4-D18 ade6-M210 h⁻* | This study |
| KGY6053-2 | *cdc15-31A rlc1-mNG:hyg$^{R}$ sid4-mNG:Kan$^{R}$ ade6-21x ura4-D18 leu1-32 h$^{+}$* | This study |

| | | |
|---|---|---|
| KGY19083 | *rlc1-mNeonGreen:hygR sid4-mNeonGreen:kan$^R$ ade6-M21X ura4-d18 leu1-32 h$^+$* | Lab stock |
| KGY1498-2 | *mNG-cdc15 ade6-M21X ura4-D18 leu1-32 h$^-$* | Lab stock |
| KGY6011-2 | *mNG-Cdc15-31A ade6-21x ura4-D18 leu1-32 h$^-$* | This study |
| **Figure 5** | | |
| KGY1498-2 | *mNG-cdc15 ade6-M21X ura4-D18 leu1-32 h$^-$* | Lab stock |
| KGY4013-2 | *pom1$^{as1}$-tdTomato:nat$^R$ mNG-cdc15 ade6-M21X ura4-D18 leu1-32 h$^-$* | This study |
| KGY16119 | *kin1$^{as1}$-FLAG:kan$^R$ mNG-cdc15 ade6-M21X ura4-D18 leu1-32 h$^+$* | This study |
| KGY1753-3 | *shk1::ClonNat$^R$ shk1$^{as2}$(M460A)-Hph$^R$ mNG-cdc15 ura4-294 leu1-32 ade6-M21X h$^-$* | This study |
| KGY1766-3 | *pck1$^{as2}$-FLAG:kan$^R$ mNG-cdc15 ura4-294 leu1-32 ade6-M21X h$^-$* | This study |
| KGY4110-2 | *pom1$^{as1}$-tdTomato:nat$^R$ kin1$^{as1}$-FLAG:kan$^R$ mNG-cdc15 ade6-M21X ura4-D18 leu1-32 h$^-$* | This study |
| KGY4075-2 | *pom1$^{as1}$-tdTomato:nat$^R$ shk1::ClonNat$^R$ shk1$^{as2}$(M460A)-Hph$^R$ mNG-cdc15 ade6-M21X ura4-D18 leu1-32 h$^-$* | This study |
| KGY4097-2 | *pom1$^{as1}$-tdTomato:nat$^R$ pck1$^{as2}$-FLAG:kan$^R$mNG-cdc15 ade6-M21X ura4-D18 leu1-32 h$^-$* | This study |
| KGY4132-2 | *kin1$^{as1}$-FLAG:kan$^R$ shk1::ClonNat$^R$ shk1$^{as2}$(M460A)-Hph$^R$ mNG-cdc15 ade6-M21X ura4-D18 leu1-32 h$^-$* | This study |
| KGY4151-2 | *kin1$^{as1}$-FLAG:kan$^R$ pck1$^{as2}$-FLAG:kan$^R$mNG-cdc15 ade6-M21X ura4-D18 leu1-32 h$^-$* | This study |
| KGY4133-2 | *shk1::ClonNat$^R$ shk1$^{as2}$(M460A)-Hph$^R$ pck1$^{as2}$-FLAG:kan$^R$mNG-cdc15 ade6-M21X ura4-D18 leu1-32 h$^-$* | This study |
| KGY4149-2 | *pom1$^{as1}$-tdTomato:nat$^R$ kin1$^{as1}$-FLAG:kan$^R$ shk1::ClonNat$^R$ shk1$^{as2}$(M460A)-Hph$^R$ mNG-cdc15 ade6-M21X ura4-D18 leu1-32 h$^-$* | This study |
| KGY4148-2 | *pom1$^{as1}$-tdTomato:nat$^R$ shk1::ClonNat$^R$ shk1$^{as2}$(M460A)-Hph$^R$pck1$^{as2}$-FLAG:kan$^R$ mNG-cdc15 ade6-M21X ura4-D18 leu1-32 h$^-$* | This study |
| KGY4152-2 | *pom1$^{as1}$-tdTomato:nat$^R$ kin1$^{as1}$-FLAG:kan$^R$ pck1$^{as2}$-FLAG:kan$^R$mNG-cdc15 ade6-M21X ura4-D18 leu1-32 h$^-$* | This study |
| KGY4153-2 | *kin1$^{as1}$-FLAG:kan$^R$ shk1::ClonNat$^R$ shk1$^{as2}$(M460A)-Hph$^R$ pck1$^{as2}$-FLAG:kan$^R$mNG-cdc15 ade6-M21X ura4-D18 leu1-32 h$^-$* | This study |
| KGY1183-3 | *mNG-Cdc15-1A ade6-21x ura4-D18 leu1-32 h$^-$* | This study |
| KGY706-2 | *mNG-Cdc15-5A ade6-21x ura4-D18 leu1-32 h$^-$* | This study |
| KGY6007-2 | *mNG-Cdc15-11A ade6-21x ura4-D18 leu1-32 h$^-$* | This study |
| KGY19616 | *mNG-Cdc15-22A ade6-21x ura4-D18 leu1-32 h$^-$* | Lab stock |
| KGY6011-2 | *mNG-Cdc15-31A ade6-21x ura4-D18 leu1-32 h$^-$* | This study |
| KGY3019 | *cdc15-GFP::kan$^R$ ade6-M210 ura4-D18 leu1-32 h$^-$* | Lab stock |
| KGY9446 | *cdc15::cdc15(SP11A)-GFP:kan$^R$ ade6-M210 ura4-D18 leu1-32 h$^-$* | Lab stock |
| KGY9444 | *cdc15::cdc15(RXXS13A)-GFP:kan$^R$ ade6-M210 ura4-D18 leu1-32 h$^-$* | Lab stock |
| KGY8461 | *cdc15::cdc15(SP11+others-18A)-GFP:kan$^R$ ade6-M210 ura4-D18 leu1-32 h$^-$* | Lab stock |
| KGY10307 | *cdc15::cdc15(RXXS+SP+others27A)-GFP:kan$^R$ ade6-M210 ura4-D18 leu1-32 h$^+$* | Lab stock |

| **Figure 9** | | |
|---|---|---|
| KGY1498-2 | *mNG-cdc15 ade6-M21X ura4-D18 leu1-32 h⁻* | Lab stock |
| KGY19616 | *mNG-Cdc15-22A ade6-21x ura4-D18 leu1-32 h⁻* | Lab stock |
| KGY6011-2 | *mNG-Cdc15-31A ade6-21x ura4-D18 leu1-32 h⁻* | This study |
| KGY6058-2 | *cdc15-mCherry:Nat$^R$ rlc1-mNG:hyg$^R$ ade6-M210 ura4-D18 leu1-32 h⁻* | This study |
| KGY6061-2 | *cdc15-31A-mCherry:Nat$^R$ rlc1-mNG:hyg$^R$ ade6-M210 ura4-D18 leu1-32 h⁻* | This study |
| KGY5994-2 | *cdc15-mCherry:Nat$^R$ fic1-mNG:Kan$^R$ ade6-M21x ura4-D18 leu1-32 h-* | This study |
| KGY5993-2 | *cdc15-31A-mCherry:Nat$^R$ fic1-mNG:Kan$^R$ ade6-M21x ura4-D18 leu1-32 h-* | This study |
| **Figure 5-figure sup1** | | |
| KGY1498-2 | *mNG-cdc15 ade6-M21X ura4-D18 leu1-32 h⁻* | Lab stock |
| KGY4013-2 | *pom1$^{as1}$-tdTomato:nat$^R$ mNG-cdc15 ade6-M21X ura4-D18 leu1-32 h⁻* | This study |
| KGY16119 | *kin1$^{as1}$-FLAG:kan$^R$ mNG-cdc15 ade6-M21X ura4-D18 leu1-32 h⁺* | This study |
| KGY1753-3 | *shk1::ClonNat$^R$ shk1$^{as2}$(M460A)-Hph$^R$ mNG-cdc15 ura4-294 leu1-32 ade6-M21X h⁻* | This study |
| KGY1766-3 | *pck1$^{as2}$-FLAG:kan$^R$ mNG-cdc15 ura4-294 leu1-32 ade6-M21X h⁻* | This study |
| KGY4110-2 | *pom1$^{as1}$-tdTomato:nat$^R$ kin1$^{as1}$-FLAG:kan$^R$ mNG-cdc15 ade6-M21X ura4-D18 leu1-32 h⁻* | This study |
| KGY4075-2 | *pom1$^{as1}$-tdTomato:nat$^R$ shk1::ClonNat$^R$ shk1$^{as2}$(M460A)-Hph$^R$ mNG-cdc15 ade6-M21X ura4-D18 leu1-32 h⁻* | This study |
| KGY4097-2 | *pom1$^{as1}$-tdTomato:nat$^R$ pck1$^{as2}$-FLAG:kan$^R$mNG-cdc15 ade6-M21X ura4-D18 leu1-32 h⁻* | This study |
| KGY4132-2 | *kin1$^{as1}$-FLAG:kan$^R$ shk1::ClonNat$^R$ shk1$^{as2}$(M460A)-Hph$^R$ mNG-cdc15 ade6-M21X ura4-D18 leu1-32 h⁻* | This study |
| KGY4151-2 | *kin1$^{as1}$-FLAG:kan$^R$ pck1$^{as2}$-FLAG:kan$^R$mNG-cdc15 ade6-M21X ura4-D18 leu1-32 h⁻* | This study |
| KGY4133-2 | *shk1::ClonNat$^R$ shk1$^{as2}$(M460A)-Hph$^R$ pck1$^{as2}$-FLAG:kan$^R$mNG-cdc15 ade6-M21X ura4-D18 leu1-32 h⁻* | This study |
| KGY4149-2 | *pom1$^{as1}$-tdTomato:nat$^R$ kin1$^{as1}$-FLAG:kan$^R$ shk1::ClonNat$^R$ shk1$^{as2}$(M460A)-Hph$^R$ mNG-cdc15 ade6-M21X ura4-D18 leu1-32 h⁻* | This study |
| KGY4148-2 | *pom1$^{as1}$-tdTomato:nat$^R$ shk1::ClonNat$^R$ shk1$^{as2}$(M460A)-Hph$^R$pck1$^{as2}$-FLAG:kan$^R$ mNG-cdc15 ade6-M21X ura4-D18 leu1-32 h⁻* | This study |
| KGY4152-2 | *pom1$^{as1}$-tdTomato:nat$^R$ kin1$^{as1}$-FLAG:kan$^R$ pck1$^{as2}$-FLAG:kan$^R$mNG-cdc15 ade6-M21X ura4-D18 leu1-32 h⁻* | This study |
| KGY4153-2 | *kin1$^{as1}$-FLAG:kan$^R$ shk1::ClonNat$^R$ shk1$^{as2}$(M460A)-Hph$^R$ pck1$^{as2}$-FLAG:kan$^R$mNG-cdc15 ade6-M21X ura4-D18 leu1-32 h⁻* | This study |
| KGY1498-2 | *mNG-cdc15 ade6-M21X ura4-D18 leu1-32 h⁻* | Lab stock |
| KGY1183-3 | *mNG-Cdc15-1A ade6-21x ura4-D18 leu1-32 h⁻* | This study |
| KGY706-2 | *mNG-Cdc15-5A ade6-21x ura4-D18 leu1-32 h⁻* | This study |
| KGY6007-2 | *mNG-Cdc15-11A ade6-21x ura4-D18 leu1-32 h⁻* | This study |
| KGY19616 | *mNG-Cdc15-22A ade6-21x ura4-D18 leu1-32 h⁻* | Lab stock |

| | | |
|---|---|---|
| KGY6011-2 | *mNG-Cdc15-31A ade6-21x ura4-D18 leu1-32 h*$^{-}$ | This study |
| **Figure 5-figure sup 2** | | |
| KGY246 | *ade6-M210 ura4-D18 leu1-32 h*$^{-}$ | Lab stock |
| KGY2711 | *mid1::ura4*$^{+}$ *ade6-M210 ura4-D18 leu1-32 h*$^{+}$ | Lab stock |
| KGY4951-2 | *mid1::ura4*$^{+}$ *pom1-as1-tdTomato:nat*$^{R}$ *ade6-M210 ura4-D18 leu1-32 h*$^{+}$ | Lab stock |
| KGY1516-2 | *kin1*$^{as1}$*-FLAG:kan*$^{R}$ *ura4-294 leu1-32 ade6-704 h*$^{+}$ | This study |
| KGY19282 | *shk1::ClonNat*$^{R}$ *shk1*$^{as2}$*(M460A)-Hph*$^{R}$ *leu1-32 ura4-D18 ade6-M210 h*$^{-}$ | (Cipak et al., 2011) |
| KGY526-2 | *pck1*$^{as2}$*-FLAG:kan*$^{R}$ *ade6-M21X leu1-32 ura4-D18 h*$^{+}$ | Lab stock |
| KGY19781 | *mid1::ura4*$^{+}$ *kin1*$^{as1}$*-FLAG:kan*$^{R}$ *ura4-294 leu1-32 ade6-704 h*$^{+}$ | This study |
| KGY19782 | *mid1::ura4*$^{+}$ *pck1*$^{as2}$*-FLAG:kan*$^{R}$ *ade6-M21X leu1-32 ura4-D18 h*$^{+}$ | This study |
| KGY5911-2 | *mid1::ura4*$^{+}$ *shk1::ClonNat*$^{R}$ *shk1*$^{as2}$*(M460A)-Hph*$^{R}$ *leu1-32 ura4-D18 ade6-M210 h*$^{-}$ | This study |
